# Physiological Differences Between Advanced Crossfit Athletes, Recreational Crossfit Participants, and Physically-Active Adults

**DOI:** 10.1101/782359

**Authors:** Gerald T. Mangine, Matthew T. Stratton, Christian G. Almeda, Michael D. Roberts, Tiffany A. Esmat, Trisha A. VanDusseldorp, Yuri Feito

## Abstract

This investigation examined anthropometric, hormonal, and physiological differences between advanced (ADV; n = 8, 27.8 ± 4.2 years, 170 ± 11 cm, 79.8 ± 13.3 kg) and recreational (REC; n = 8, 33.5 ± 8.1 years, 172 ± 14 cm, 76.3 ± 19.5 kg) CrossFit (CF) trained participants in comparison to physically-active controls (CON; n = 7, 27.5 ± 6.7 years, 171 ± 14 cm, 74.5 ± 14.3 kg). ADV and REC were distinguished by their past competitive success. REC and CON were resistance-trained (>2 years) and exercised on 3-5 days·wk^-1^ for the past year, but CON utilized traditional resistance and cardiovascular exercise. All participants provided a fasted, resting blood sample and completed assessments of resting metabolic rate, body composition, muscle morphology, isometric mid-thigh pull strength, peak aerobic capacity, and a 3-minute maximal cycle ergometer sprint across two separate occasions (separated by 3-7 days). Blood samples were analyzed for testosterone, cortisol, and insulin-like growth factor-1. One-way analysis of variance revealed ADV to possess lower body fat percentage (6.7-8.3%, *p* = 0.007), greater bone and non-bone lean mass (12.5-26.8%, *p* ≤ 0.028), muscle morphology characteristics (14.2-59.9%, *p* < 0.05), isometric strength characteristics (15.4-41.8%, *p* < 0.05), peak aerobic capacity (18.8-19.1%, *p* = 0.002), and anaerobic performance (15.4-51.1%, *p* ≤ 0.023) compared to both REC and CON. No differences were seen between REC and CON, or between all groups for resting metabolic rate or hormone concentrations. These data suggest ADV possess several physiological advantages over REC and CON, whereas similar physiological characteristics were present in individuals who have been regularly participating in either CF or resistance and cardiovascular training for the past year.

## INTRODUCTION

CrossFit® (CF) is a form of high-intensity functional training that combines resistance exercises, gymnastics, and traditional aerobic modalities (e.g., cycling, rowing, running) into single workouts that vary by day to elicit general physical preparedness (1, 2). This training form is enjoyed recreationally by participants of varying levels of fitness, training experience, age, and lifestyles (3) and also exists as its own sport. The primary CF competition is the Reebok CrossFit Games™ (the Games) which awards individual winners the title of “Fittest on Earth™”. Historically, this competition has consisted of several stages designed to narrow the initial participant pool (>400,000 athletes) down to the top athletes within each category (i.e., adult men, adult women, teenagers, Masters, teams). Although the existence, format and relevance of each stage have undergone changes throughout the competition’s existence (4, 5), the presence of an initial online qualifying round (e.g., the CrossFit Open^TM^) has remained. This round typically involves multiple workout challenges that are completed over the course of several weeks. Competitors who complete all workouts and rank high enough will progress to the next stage of the competition (i.e., historically regionals, currently the Games or Sanctioned competitions that lead to the Games). Regardless of which stage, it is expected that each workout will consist of a set of challenges that will require some combination of strength, power, endurance, and/or sport-specific skill (1). However, little is known about which physiological factors are most influential of CF competition performance or distinguish competitive status.

Body mass (6), strength and anaerobic power (6–10), aerobic capacity (9), sport-specific skill (8, 10), and experience (9) have all been associated with either workout performance or competitive ranking. Collectively, these data imply that athletes must train to be proficient in each to perform well in competition. However, several limitations exist among these studies that prevent making such a conclusion. For instance, Serafini et al. (2018) reported that higher ranking competitors of the 2016 Open were stronger, more powerful, and more proficient at short-duration, sprint-type CF workouts. Among regional competitors, final ranking was positively related to 400-m sprint time and time-to-completion in longer, benchmark workouts (i.e., Filthy-50) (*r* = 0.69 – 0.77), and negatively related to maximal weight lifted in the Olympic lifts (*r* = −0.39 to −0.42) (10). Although these studies involved participants who have successful competitive records, the measures used to distinguish rank were all self-reported. As such, the authenticity and actual data of measurement (self-reported data were obtained from an online resource) cannot be verified. In contrast, others have measured a variety of physical parameters and related them to CF-style workouts performed in a controlled, laboratory setting (6, 7, 9). While these studies have also included successful CF athletes, laboratory workouts do not adequately emulate the competitive setting and may influence the physiological response to CF training (11–14). Thus, questions remain about the distinguishing characteristics of successful CF athletes.

In more traditional sports (e.g., football, baseball, basketball, etc.), identifying the key physiological and athletic characteristics that distinguish performance is common (15–18). The practice enables strength and conditioning professionals to develop sport-specific training programs that are more effective in translating adaptations to in-game performance. However, CF is unique in that typical training session workouts mirror those that appear in competition. Moreover and consistent with its primary purpose (1, 2), chronic participation in CF training has been documented to improve a variety of fitness parameters (19). Though it might be assumed that CF training represents an ideal training strategy for developing the physiological characteristics that distinguish performance in the sport, such a conclusion would be premature based on the available data.

Evidence of CF training being more advantageous towards developing a variety of fitness outcomes in comparison to alternative training strategies (e.g., resistance training, high-intensity interval training) is equivocal (19–25). This is likely because most comparative training studies have utilized untrained or novice (to CF) participants, which is problematic because they do not require a very specific or intense training stimulus to elicit adaptations compared to experienced trainees (26). It is possible that either a longer training duration or more advanced participants are necessary to observe the advantages or disadvantages of the CF strategy. Unfortunately, elite competitors rarely share their training strategies and anecdotal evidence suggests that they incorporate more than what commonly occurs during a typical CF training session. To the best of our knowledge, only one well-controlled study exists where a variety of physiological parameters were examined in trained participants (27). In that cross-sectional investigation, men with at least on year of CF training experience outperformed their resistance-trained (> 1 year) counterparts in a multi-stage shuttle run test and possessed a higher aerobic capacity; all other measures were statistically similar. While this study provides evidence in favor of CF training, there was no aerobic training requirement for the resistance-trained group, and the actual experience of the CF group was unclear beyond their having participated in the strategy for at least one year. It is possible that multiple physiological differences exist when experience is considered. Therefore, the purpose of this study was to examine anthropometric, hormonal, and physiological differences between advanced CF athletes, recreational CF practitioners, and physically-active adults who regularly participate in both resistance and cardiovascular training. Since adaptations are specific to the training modality and effort (26), we hypothesized that body composition, muscle morphology, aerobic and anaerobic performance, and strength would be different between groups. Specifically, the advanced CF athletes would outperform the other groups whereas recreational CF practitioners and physically-active adults would be similar. However, because resting hormonal concentrations do not typically change through training (14), it was hypothesized that these would be similar between groups.

## MATERIALS AND METHODS

### Experimental Design

For this cross-sectional study, physically-active adults were recruited and assigned into groups based on their experience with CF training and performance during specific CF competitions. Participants who possessed CF training experience (> 2 years) were classified as advanced (ADV) if they had previously qualified for the regional round of the Games competition. Otherwise, they were classified as recreational (REC) because they had never progressed beyond the opening round of the competition (i.e., The Open) but still trained on 3 – 5 days per week for at least the previous year. Individuals who did not possess CF training experience but possessed resistance training experience (> 2 years) and participated in both resistance and cardiovascular training on 3 – 5 days per week for at least the previous year, were assigned to the physically-active control (CON) group. All participants reported to the Exercise Physiology Laboratory on two separate occasions, within one month of the onset of the Open, to complete all testing. During the first visit, each participant provided a fasted blood sample before completing assessments of muscle morphology and peak aerobic capacity. Participants returned to the Exercise Physiology Lab for the second visit (within 3 – 7 days of the first visit) to complete assessments of resting metabolic rate, body composition, strength, and anaerobic performance. All testing sessions occurred in the morning (∼6:00 – 10:00 a.m.) with the participants having abstained from unaccustomed physical activity and alcohol for 24 hours, caffeine for 12 hours, and fasted for 8 hours. Participants completed all measurements while wearing comfortable athletic clothing and were able to consume a light snack prior to performance testing (i.e., peak aerobic capacity, strength, and anaerobic performance). Prior to leaving the laboratory on the first visit, participants were asked to complete a 24-hour dietary recall, retain a copy, and follow a similar diet prior to their second visit. Comparisons were made between groups for all anthropometric, biochemical, and physiological measures.

### Participants

Twenty-three physically-active adults (29.7 ± 6.8 years, 171 ± 12 cm, 76.9 ± 15.4 kg) agreed to participate in this study. All participants were free of any physical limitations (determined by medical and physical-activity history questionnaire and PAR-Q+) and had been regularly participating (at the time of recruitment) in their chosen exercise form (i.e., CrossFit training or Resistance/Cardiovascular training) for a minimum of 2 years. Participants in ADV (n = 8, 27.8 ± 4.2 years, 170 ± 11 cm, 79.8 ± 13.3 kg) reported having regularly participated in resistance training for 11.5 ± 5.8 years and CF training for 6.4 ± 5.6 years (6 – 7 sessions·week^-1^). As individual competitors, the highest rank these participants ever achieved in the Open was 659^th^ ± 991^st^ (range: 19^th^ – 3,052^nd^) within their respective divisions worldwide. While each of these athletes qualified for this study by having competed as members of a team in regional (highest average rank = 11^th^ ± 13^th^) and Games competition (highest average rank = 20^th^ ± 9^th^), three competed individually in their respective regions with one having progressed to the Games on multiple occasions. REC participants (n = 8, 33.5 ± 8.1 years, 172 ± 14 cm, 76.3 ± 19.5 kg) reported having regularly participated in resistance training for 8.1 ± 7.9 years and CF training for 3.3 ± 1.7 years (4 – 5 sessions·week^-1^). The highest rank these participants had ever achieved in the Open was 22,306th ± 14,028^th^ (range: 5,466th – 44,315^th^) within their respective divisions worldwide. Participants in CON (n = 7, 27.5 ± 6.7 years, 171 ± 14 cm, 74.5 ± 14.3 kg) reported having 7.6 ± 4.8 years of regular resistance training experience and incorporated 3.7 ± 1.3 sessions and 3.6 ± 1.0 sessions of resistance and cardiovascular training per week. Although two participants in CON reported having previously participated in CF-style workouts, these did not occur with regularity (< 3 sessions·week^-1^) or for an extended duration (< 1 year) and they had never competed in the Open at the time of data collection. Following an explanation of all procedures, risks and benefits, each participant provided his or her written informed consent to participate in the study. The study was conducted in accordance with the Declaration of Helsinki, and the protocol was approved by the Kennesaw State University Institutional Review Board (#17-501).

### Blood sampling and biochemical analysis

Blood samples were obtained on the first visit prior to any physical activity. All samples were obtained from an antecubital vein using a needle by a research team member who was trained and experienced in phlebotomy. Approximately 15 mL of blood was drawn into SST tubes (for serum collection) and EDTA-treated Vacutainer® tubes (for plasma). SST tubes were allowed to clot for 10 minutes prior to centrifugation, while EDTA treated tubes were centrifuged immediately for 10 minutes at 3600 rpms at 4 °C. The resulting serum and plasma were aliquoted and stored at −80°C until analysis.

Circulating concentrations of testosterone (T; in ng·dL^-1^), cortisol (C; in μg·dL^-1^), and insulin-like growth factor (IGF-1; in ng·mL^-1^) were assessed via enzyme-linked immunosorbent assays (ELISA) via a 96-well spectrophotometer (BioTek, Winooski, VT) using commercially available kits. To eliminate inter-assay variance, all samples for each assay were thawed once and analyzed in duplicate in the same assay run by a single technician. Samples were analyzed in duplicate, with an average coefficient of variation of 1.63% for T, 6.88% for C, and 2.00% IGF-1.

### Muscle morphology

Non-invasive skeletal muscle ultrasound images were collected from the right thigh and arm locations of all participants. This technique uses sound waves at fixed frequencies to create *in vivo*, real time images of the limb musculature. Prior to image collection, all anatomical locations of interest were identified using standardized landmarks for the rectus femoris (RF), vastus medialis (VM), vastus lateralis (VL), biceps brachii (BB), and triceps brachii (TB) muscles. The landmarks for the thigh musculature were identified along the longitudinal distance over the femur. The RF and VM were respectively assessed at 50% and 20% of the distance from the proximal border of the patella to the anterior, inferior suprailiac crest. The VL was assessed at 50% of the distance from the lateral condyle of the tibia to the most prominent point of the greater trochanter of the femur. VL measurement required the participant to lay on their side. Landmark identification of the BB and TB required the participant to sit upright on the examination table and extend their arm to rest upon the shoulder of the researcher. Both muscles were assessed along the humerus at a position equal to 40% of the distance from the lateral epicondyle to the acromion process of the scapula (28). Subsequently, the participant resumed laying supine on the examination table for a minimum of 5 – 10 minutes to allow fluid shifts to occur before images were collected (29). The same investigator performed all landmark measurements for each participant.

A 12 MHz linear probe scanning head (General Electric LOGIQ S7 Expert, Wauwatosa, WI, USA) was coated with water soluble transmission gel to optimize spatial resolution and used to collect all ultrasound images. Collection of each image began with the probe being positioned on (and perpendicular to) the surface of the skin to provide acoustic contact without depressing the dermal layer. Subsequently, two consecutive images were collected in the extended field of view mode (Gain = 50 dB; Image Depth = 5 – 6 cm) using a cross-sectional sweep in the axial plane to capture panoramic images of each muscle. At the same sites, two consecutive images were collected with the probe oriented longitudinal to the muscle tissue interface using Brightness Mode (B-mode) ultrasound (30). Each of these images included a horizontal line (approximately 1 cm), located below the image, which was used for calibration purposes when analyzing the images offline (31). To capture images of the RF and VM, the participant remained in the supine position, with their legs extended but relaxed. A rolled towel was placed beneath the popliteal fossa of the dominant leg, allowing for a 10° bend in the knee as measured by a goniometer, and the dominant foot secured (32). For the VL, the participant was placed on their side with their legs together and the rolled towel between their needs. Once again, the legs were positioned to allow a 10° bend in the knees, as measured by a goniometer (32). Measurement of the BB and TB required the participant to sit upright with their arm extended, resting on the shoulder of the researcher. The same investigator positioned each participant and collected all images.

After all images were collected, the ultrasound data were transferred to a personal computer for analysis via Image J (National Institutes of Health, Bethesda, MD, USA, version 1.45s) by the same technician. All panoramic images were used to measure cross-sectional area (CSA) and echo intensity. For these measures, the polygon tracking tool in the ImageJ software was used to isolate as much lean muscle as possible without any surrounding bone or fascia (30). Subsequently, Image J calculated the area contained within the traced muscular image and reported this value in centimeters squared (± 0.1cm^2^). Concurrently, echo intensity was determined by grayscale analysis using the standard histogram function in ImageJ (30) and expressed as an arbitrary unit (au) value between 0 – 255 (0: black; 255: white) with lower values reflecting more contractile tissue within each muscle (30, 33). Mean echo intensity values were then corrected for subcutaneous fat thickness (SFT; averaged from the SFT values obtained at the medial, midline, and lateral sites of each muscle) using Equation 1 (34). All B-mode images were used to measure muscle thickness (± 0.01 cm; perpendicular distance between the superficial and deep aponeuroses) and pennation angle (± 0.1°; intersection of the fascicles with the deep aponeurosis). Fascicle length (± 0.1 cm) across the deep and superficial aponeuroses was estimated from muscle thickness and pennation angle using Equation 2. Intraclass correlation coefficients (ICC_3,k_ = 0.77 – 0.99) for determining muscle thickness, pennation angle, CSA and echo intensity was previously determined in ten active, resistance-trained men (25.3 ± 2.0 years, 180 ± 7 cm, 90.8 ± 6.8 kg) using the methodology described above. The methodology for determination of FL has a reported estimated coefficient of variation of 4.7% (35).

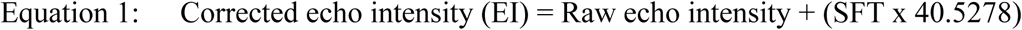

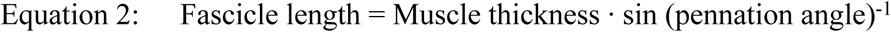

### Peak aerobic capacity

Peak aerobic capacity (VO_2_peak; ml·kg^-1^·min^-1^), respiratory compensation threshold (RCT; ml·kg^-1^·min^-1^), and gas exchange threshold (GET; ml·kg^-1^·min^-1^) were assessed using a continuous, ramp exercise protocol performed on an electromagnetic-braked cycle ergometer (Lode Excalibur Sport, Lode., B.V., Groningen, The Netherlands). Prior to testing, each participant completed a standardized warm-up that consisted of riding a cycle ergometer for 5 minutes at the participant’s preferred resistance and cadence followed by 10 body weight squats, 10 alternating lunges, 10 walking knee hugs and 10 walking butt kicks. Participants were then permitted to continue their warm-up with any additional practices that would help them feel comfortable entering the test. Participants were fitted with a heart rate (HR) monitor (Team^2^, Polar, Lake Success, NY), a nose clip, and a 2-way valve mask connected to a metabolic measurement system (True One 2400, ParvoMedics Inc., Salt Lake City, UT) to measure expired gases. The cycle ergometer seat height and handlebar distance were adjusted to the participant’s comfort. The participants initially completed a 3-minute warm-up period with the resistance set at 50 W before starting the test at 75 W. During testing, the participants were asked to maintain a self-selected pedaling rate (> 50 rpm’s) while power output was increased by 25 W every minute until volitional fatigue or pedaling rate dropped below 50 rpm’s for longer than 15 seconds. Upon completion of the test, each participant immediately progressed to a 3-minute active recovery period where they continued to pedal at their own cadence against a 50 W load. HR was assessed on each minute of the 3-minute recovery period. Participants were then removed from the cycle ergometer and asked to rest in a chair for an additional two minutes.

Relative oxygen consumption values (i.e., VO_2_·kg^-1^) collected on each breath were averaged using the 11-breath averaging technique (36) and used to determine the highest value achieved during the test (i.e., VO_2_peak). RCT, also known as the second ventilatory threshold, was identified as the VO_2_ value at which the increase in ventilation-VO_2_ relationship was accompanied by an increase in the ventilation-VCO_2_ relationships (37). The GET was determined using the V-slope method described by Beaver et al. (38). The GET was defined as the VO_2_ value corresponding to the intersection of two linear regression lines derived separately from the data points below and above the breakpoint in the CO_2_ produced (VCO_2_) versus the VO_2_ relationship (39).

### Dietary recall

Participant’s dietary intake was tracked for the 24-hour period preceding each visit via a paper dietary food recall form. All participants were instructed on how to properly log their food, snacks and drinks via the paper form. Specifically, following their enrollment on their first visit, participants were asked to record their food intake (breakfast, lunch, dinner, drinks and snacks) for the previous 24 hours prior. Prior to leaving the laboratory on the first visit, the participants were given a copy of their food recall form and asked to consume a similar diet during the 24 hours prior to their second visit. Each form was visually inspected to confirm dietary compliance.

### Resting metabolic rate assessment

Resting metabolic rate (RMR, kcals·day^-1^) assessment was completed in a quiet room with minimal lighting (e.g., only light from the RMR machine) located within the Exercise Physiology Laboratory. Prior to their arrival, participants were informed of all pre-test guidelines as outlined by Compher et al. (40). These included: 1) avoiding alcohol consumption 24 hours prior to testing, 2) no food or caffeine ingestion 8 and 12 hours prior to testing, respectively, and 3) discontinuing unaccustomed physical activity 24 hours prior to testing. Resting metabolic rate was measured via a metabolic measurement system (Parvo Medics TrueOne 2400, ParvoMedics Inc., Salt Lake City, UT) utilizing a ventilated hood. Participants were asked to rest in the supine position with the ventilated hood placed over their face and neck for a maximum of 30 minutes. RMR determination was based on a 5-minute interval of measured volume of oxygen consumption (VO_2_) with a coefficient of variation less than 10% (40). The average coefficient of variation was 6.36%.

### Body composition assessments

Initially, height (± 0.1 cm) and body mass (± 0.1 kg) were determined using a stadiometer (WB-3000, TANITA Corporation, Tokyo, Japan) with the participants standing barefoot, with feet together, in their normal daily attire. Subsequently, body composition was assessed by three common methods (i.e., dual energy X-ray absorptiometry [iDXA, Lunar Corporation, Madison, WI], air displacement plethysmography [BodPod, COSMED USA Inc., Chicago, IL], and bioelectrical impedance analysis [770 Body Composition and Body Water Analyzer, InBody, Seoul, South Korea]) using standardized procedures. Briefly, iDXA scanning required participants to remove any metal or jewelry and lay supine on the iDXA table prior to an entire body scan in “standard” mode using the company’s recommended procedures and supplied algorithms. Quality assurance was assessed by daily calibrations performed prior to all scans using a calibration block provided by the manufacturer. All iDXA measurements were performed by the same researcher using standardized subject positioning procedures. For air displacement plethysmography, the device and associated scale were calibrated daily using a known volume and mass provided by the manufacturer. During testing, participants were asked to wear a tight-fitting bathing suit or compression shorts and swim cap before entering the device. Two trials were performed for each participant to obtain two measurements of body volume within 150 mL. A third trial was performed if body volume estimates from the first two trials were not within 150 mL, and values from the two closest trials were averaged. Thoracic lung volume was estimated (41). Bioelectrical impedance analysis required participants to stand barefoot on two metal sensors located at the base of the device and hold two hand grips for approximately 30 – 60 seconds. Prior to stepping onto the device, participants cleaned the soles of their feet with alcohol wipes provided by the manufacturer.

Following testing, body mass, bone mineral content (BMC; from iDXA), body volume (from BodPod), and total body water (from bioelectrical impedance analysis) were entered into a 4-compartment model, Equation 3 to estimate body fat percentage (BF%) (42), fat mass (± 0.1 kg), and fat-free mass (± 0.1 kg). These values, along with regional (arms [sum of each arm], legs [sum of each leg], and trunk [sum of spine and pelvis]) estimates of bone mineral content (±0.1 kg) and non-bone lean mass (± 0.1 kg) obtained from iDXA following manual demarcation of these regions of interest were used for all group comparisons. Intraclass correlation coefficients (ICC_3,1_ = 0.74 – 0.99) for manually determining regional estimates of bone mineral content and non-bone lean mass had been previously found in 10 healthy, physically-active adults (25.1 ± 2.4 years; 176 ± 7 cm, 81.1 ± 18.5 kg).

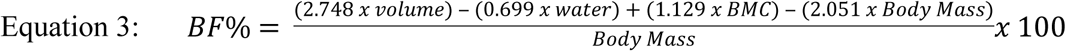

### Strength assessment

Following RMR and body composition assessments, strength was assessed by an isometric mid-thigh pull test. Prior to testing, each participant completed the same standardized warm-up described for the first visit (i.e., 5 minutes of cycling, dynamic stretching, additional self-selected warm-up practices) followed by a protocol specific to the isometric mid-thigh pull test. The specific component included three isometric efforts on an immobilized barbell positioned at approximately the mid-thigh using a perceived intensity of 50, 70, and 90% of maximum effort, interspersed with a one-minute recovery. The specific warm-up and isometric mid-thigh pull test were completed within a power rack (Rogue Fitness, Columbus, OH) while standing upon a portable force plate (Accupower, AMTI, Watertown, MA). While standing on the force plate, the mid-thigh position was determined for each participant before testing by marking the midpoint distance between the knee and hip joints. Each participant was instructed to assume their preferred second pull power-clean position by self-selecting their hip and knee angles. The height of the barbell was adjusted to a position approximately equal (± 2.54 cm) to the mid-thigh. The participants were then asked to use an overhand, hooked grip on the barbell. The hook grip was selected for this test because all participants reported having had experience with the technique and it is commonly used among CF athletes during competition. Participants were also allowed to wrap their thumbs with athletic training tape and use chalk. Upon the researcher’s “3, 2, 1, Go!” command, the participants were instructed to pull upwards on the barbell as hard and as fast as possible and to continue their maximal effort for 6 seconds. All participants were instructed to relax before the command “GO!” to avoid precontraction and were allotted three maximal attempts. The portable force plate measured the ground reaction forces, imposed onto the plate by the participant, as he/she pulled upon the bar. Peak force (F; in N) production, peak and average rate of force development (RFD_PEAK_, RFD_AVG_; in N·s^-1^), and F and RFD across specific time bands (i.e., 0–30, 0–50, 0–90, 0–100, 0–150, 0–200, and 0–250 milliseconds) were subsequently calculated, as previously described (43).

### Anaerobic performance assessment

Following the strength assessment, anaerobic performance was assessed via a 3-minute maximal sprint on an electromagnetic-braked cycle ergometer (Lode Excalibur Sport, Lode., B.V., Groningen, The Netherlands). Prior to the test, seat height and handlebar positions were adjusted to mirror their positions during the peak aerobic capacity test, and participants were provided with time (∼3 – 5 minutes) to acclimate to the cycle ergometer. A 5-minute rest period was then allotted before initiating the testing protocol, which has been previously described in detail elsewhere (44). Briefly, the test began with a 1-minute baseline period that involved 55 seconds of unloaded cycling at 90 rpm and then accelerating up to approximately 110 rpm over the last 5 seconds of the minute. The protocol immediately transitioned to the 3-minute testing period where the participants attempted to maintain cadence as high as possible throughout its entirety. Resistance for the test was set using the linear mode of the cycle ergometer (linear factor = power / [preferred cadence]^2^). That is, the linear factor was calculated as the power output halfway between the VO_2_peak and GET, divided by the preferred cadence of untrained cyclists (70 rpm^2^) (45–47). To prevent pacing and ensure an all-out effort, participants were not informed of the elapsed time and strong verbal encouragement was provided. After 3 minutes, the participants progressed to a 3-minute recovery stage at 50 Watts at their preferred cadence. Peak power (± 1 W), critical power (CP; average power over the final 30 seconds of the test; ± 1 W) (46), and anaerobic work capacity (AWC; work done above CP; (± 0.1 kJ) (47) were calculated based upon performance during the 3-minute sprint test.

### Statistical analysis

Data were modeled using both a frequentist and Bayesian approach. The frequentist approach involved a two-way (Group x Sex) analysis of variance (ANOVA) for each dependent variable. Assumptions of normality and equal variance were verified by Shapiro-Wilk and Levene’s tests, respectively. Significant interactions and main effects were further examined using Tukey’s post-hoc analysis. Criterion alpha was set at *p* ≤ 0.05. To further assess the likelihood (or the effect of group and/or sex) of the data under the alternative hypothesis compared to the null hypothesis, a two-way Bayesian ANOVA was performed with default prior scales (48). Likelihood was represented in the form of Bayes factors (i.e., BF_10_) and were interpreted according to the recommendations of Wagenmakers et al. (49). That is, data were interpreted as evidence in favor of the null hypothesis when BF_10_ < 1. Otherwise, it was interpreted as “anecdotally” (1 < BF_10_ < 3), “moderately” (3 < BF_10_ < 10), “strongly” (10 < BF_10_ < 30), “very strongly” (30 < BF_10_ < 100), or “extremely” (BF_10_ > 100) in favor of the alternative hypothesis. All statistical analyses were performed using JASP 0.10.2 (Amsterdam, the Netherlands). All data are reported as mean ± standard deviation.

## RESULTS

### Resting hormone concentrations

No interactions were observed for T, C, IGF-1. However, a trend for an interaction (F = 2.87, *p* = 0.090) driven by a main sex effect was seen for T (F = 6.11, *p* = 0.027) with anecdotal differences between sexes being 2.058 times likely compared to the null hypothesis. Specifically, women in ADV tended to exhibit lower T concentrations (*p* = 0.83) than ADV men. Male and female hormone concentrations are illustrated in Fig 1.

**Fig 1.**
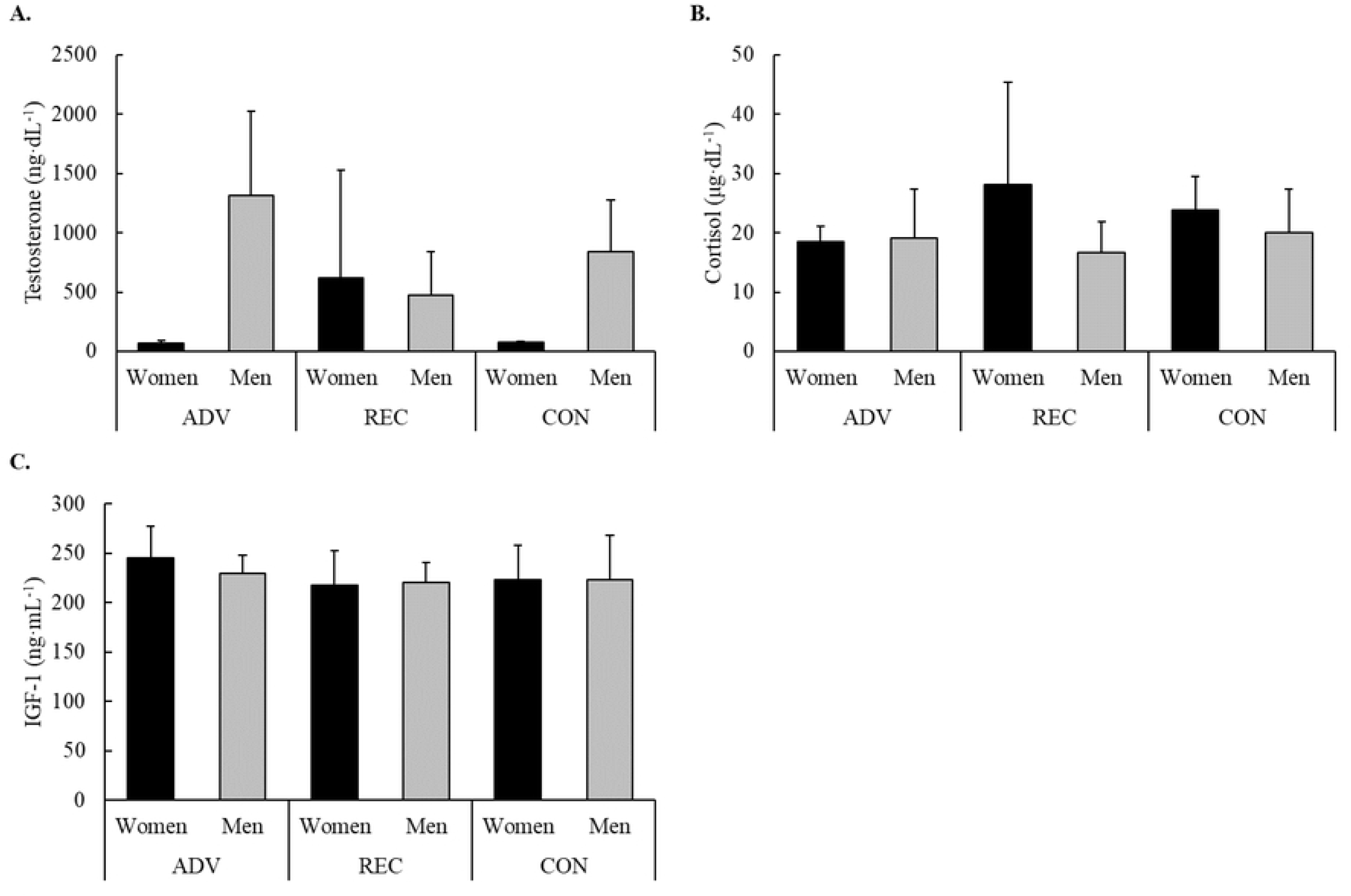
Male and female resting concentrations in A) testosterone, B) cortisol, and C) IGF-1.

### Muscle morphology

Measures of muscle morphology for each group and sex are presented in Table 1. Significant (*p* < 0.05) group x sex interactions were observed for BB fascicle length and EI for each muscle, though the likelihood of these interactions favored the null hypotheses (BF_10_ < 1). Rather, the observed interactions were primarily driven by *anecdotal*-to-*strong* evidence (1.7 < BF_10_ < 30.0) of main effects for sex and group. The observed interaction for BB fascicle length was primarily driven by a main effect for sex where women were 8.8 times more likely to possess shorter fascicles than men, specifically REC women compared to the men of REC (*p* = 0.029) and CON (*p* = 0.012). Though the underlying causes for the interactions seen for EI varied with each muscle, *anecdotal*-to-*moderate* evidence indicated that men were 1.7 – 5.5 times more likely to possess a lower EI than women. Specifically, women in REC possessed higher EI (p < 0.05) than men in ADV (RF, VL, and TB; a trend [*p* = 0.056] for VM) and REC (RF, VM, VL, and TB; a trend [*p* = 0.087] was noted for BB), and tended (p < 0.10) to be higher than men in CON (RF, VL, and TB). Even though a main effect was not seen, the effect of group was 2.4 – 30.0 times likely to influence EI. Specifically, post-hoc analysis of the interaction showed that women in REC possessed higher EI than their counterparts in ADV (RF, VM, VL, and TB).

**Table 1.** Measures of muscle morphology by group and sex.

|  |  | ADV |  | REC |  | CON |  | Group |  | Sex |  | Group x Sex |  |
| --- | --- | --- | --- | --- | --- | --- | --- | --- | --- | --- | --- | --- | --- |
|  |  | Women | Men | Women | Men | Women | Men | *p* | BF<sub>10</sub> | *p* | BF<sub>10</sub> | *p* | BF<sub>10</sub> |
| Muscle thickness (cm) | Rectus femoris | 2.48 ± 0.36 | 3.28 ± 0.50 | 2.32 ± 0.27 | 2.86 ± 0.20 | 2.46 ± 0.25 | 2.61 ± 0.43 | 0.155 | 8.2 | 0.004 | 6.7 | 0.248 | 0.5 |
|  | Vastus medialis | 3.44 ± 0.84 | 4.35 ± 0.58 | 2.77 ± 0.08 | 4.28 ± 0.61 | 3.41 ± 0.62 | 4.26 ± 0.22 | 0.439 | >100 | 0.001 | 39.8 | 0.502 | 0.3 |
|  | Vastus lateralis | 1.92 ± 0.49 | 2.47 ± 0.39 | 1.49 ± 0.25 | 1.97 ± 0.19 | 1.63 ± 0.19 | 1.67 ± 0.28 | 0.009 | 11.9 | 0.017 | 7.9 | 0.288 | 1.7 |
|  | Biceps brachii | 3.32 ± 0.60 | 4.29 ± 0.87 | 2.62 ± 0.19 | 3.74 ± 0.71 | 2.43 ± 0.08 | 3.31 ± 0.17 | 0.013 | >100 | 0.001 | 40.6 | 0.910 | 1.3 |
|  | Triceps brachii | 2.51 ± 0.45 | 3.05 ± 0.76 | 2.44 ± 0.64 | 2.91 ± 0.58 | 1.98 ± 0.24 | 3.05 ± 0.43 | 0.674 | 7.1 | 0.008 | 2.1 | 0.541 | 0.3 |
| Pennation angle (°) | Rectus femoris | 12.7 ± 3.5 | 15.5 ± 0.5 | 10.0 ± 3.0 | 14.5 ± 1.2 | 13.3 ± 5.6 | 16.4 ± 5.5 | 0.375 | 2.9 | 0.036 | 1.4 | 0.894 | 0.5 |
|  | Vastus medialis | 19.2 ± 3.9 | 26.2 ± 9.3 | 17.2 ± 3.4 | 24.9 ± 6.0 | 27.7 ± 10.8 | 24.0 ± 3.5 | 0.417 | 0.7 | 0.216 | 0.4 | 0.229 | 0.2 |
|  | Vastus lateralis | 14.3 ± 3.1 | 14.8 ± 2.9 | 10.7 ± 3.4 | 12.4 ± 5.8 | 12.3 ± 0.4 | 13.2 ± 3.5 | 0.286 | 0.6 | 0.502 | 0.5 | 0.947 | 0.1 |
|  | Biceps brachii | 13.6 ± 3.1 | 17.5 ± 2.2 | 12.8 ± 2.1 | 12.3 ± 5.0 | 10.5 ± 2.2 | 9.7 ± 1.5 | 0.009 | 7.0 | 0.489 | 3.2 | 0.249 | 0.4 |
|  | Triceps brachii | 17.9 ± 5.1 | 26.8 ± 7.8 | 11.1 ± 3.2 | 16.8 ± 4.5 | 14.2 ± 2.6 | 20.3 ± 4.1 | 0.012 | 34.5 | 0.004 | 13.8 | 0.791 | 2.4 |
| Fascicle length (cm) | Rectus femoris | 12.2 ± 5.1 | 12.3 ± 2.2 | 14.1 ± 3.8 | 11.5 ± 0.8 | 12.8 ± 7.6 | 10.3 ± 4.5 | 0.852 | 0.6 | 0.365 | 0.3 | 0.786 | 0.1 |
|  | Vastus medialis | 10.6 ± 2.4 | 11.2 ± 4.8 | 9.6 ± 1.8 | 10.6 ± 2.7 | 7.9 ± 2.4 | 10.6 ± 1.1 | 0.552 | 0.6 | 0.280 | 0.3 | 0.758 | 0.1 |
|  | Vastus lateralis | 7.7 ± 0.4 | 9.9 ± 2.1 | 8.3 ± 1.5 | 10.5 ± 4.6 | 7.6 ± 0.6 | 7.7 ± 2.2 | 0.399 | 0.8 | 0.163 | 0.4 | 0.643 | 0.2 |
|  | Biceps brachii | 14.5 ± 2.8 | 14.2 ± 1.3 | 12.0 ± 1.1<sup>df</sup> | 19.0 ± 4.8<sup>c</sup> | 13.8 ± 2.8 | 19.9 ± 2.7<sup>c</sup> | 0.271 | 9.8 | 0.002 | 8.8 | 0.043 | 0.5 |
|  | Triceps brachii | 8.7 ± 2.8 | 7.3 ± 2.9 | 12.8 ± 1.7 | 10.5 ± 3.1 | 8.2 ± 0.9 | 9.1 ± 2.2 | 0.018 | 4.3 | 0.370 | 2.4 | 0.448 | 0.5 |
| Cross-sectional area (cm<sup>2</sup>) | Rectus femoris | 10.8 ± 2.4 | 17.8 ± 3.3 | 8.8 ± 1.5 | 15.6 ± 1.8 | 11.2 ± 1.3 | 15.0 ± 2.8 | 0.216 | >100 | 0.000 | >100 | 0.364 | 0.4 |
|  | Vastus medialis | 24.3 ± 6.6 | 29.8 ± 4 | 17.0 ± 3.9 | 27.8 ± 6.4 | 18.2 ± 2.1 | 23.7 ± 1.2 | 0.046 | 22.3 | 0.002 | 12.0 | 0.441 | 0.8 |
|  | Vastus lateralis | 29.8 ± 4.1 | 44.9 ± 4.1 | 24.4 ± 2.6 | 38.2 ± 4.5 | 24.1 ± 1.7 | 37.9 ± 3.0 | 0.004 | >100 | 0.001 | >100 | 0.912 | 0.5 |
|  | Biceps brachii | 8.4 ± 1.7 | 17.7 ± 9.2 | 7.4 ± 2.0 | 14.9 ± 2.5 | 7.9 ± 0.2 | 12.9 ± 1.0 | 0.464 | 77.5 | 0.001 | 32.6 | 0.622 | 0.3 |
|  | Triceps brachii | 10.5 ± 1.2 | 18 ± 4.3 | 7.0 ± 1.4 | 17.0 ± 5.7 | 8.9 ± 1.4 | 14.0 ± 3.6 | 0.273 | >100 | 0.000 | >100 | 0.417 | 0.3 |
| Echo intensity (au) | Rectus femoris | 116 ± 26<sup>c</sup> | 113 ± 11<sup>c</sup> | 174 ± 33<sup>abd</sup> | 97 ± 14<sup>ce</sup> | 151 ± 16<sup>d</sup> | 129 ± 14 | 0.061 | 30.0 | 0.001 | 5.5 | 0.008 | 0.6 |
|  | Vastus medialis | 105 ± 16<sup>c</sup> | 108 ± 11 | 153 ± 39<sup>ad</sup> | 104 ± 12<sup>c</sup> | 116 ± 11 | 134 ± 13 | 0.093 | 2.4 | 0.289 | 0.8 | 0.012 | 0.4 |
|  | Vastus lateralis | 111 ± 20<sup>c</sup> | 113 ± 15<sup>c</sup> | 171 ± 42<sup>abd</sup> | 107 ± 19<sup>c</sup> | 134 ± 16 | 123 ± 10 | 0.087 | 4.3 | 0.023 | 1.7 | 0.028 | 0.7 |
|  | Biceps brachii | 123 ± 26 | 140 ± 9 | 170 ± 45 | 115 ± 29 | 148 ± 9 | 142 ± 17 | 0.585 | 0.6 | 0.206 | 0.5 | 0.044 | 0.2 |
|  | Triceps brachii | 83 ± 17<sup>c</sup> | 89 ± 18<sup>c</sup> | 145 ± 43<sup>abd</sup> | 78 ± 13<sup>c</sup> | 114 ± 16 | 100 ± 6 | 0.079 | 7.2 | 0.014 | 1.8 | 0.012 | 0.6 |

Note: <sup>a</sup> = Significantly ( $p < 0.05$ ) different from ADV women; <sup>b</sup> = Significantly ( $p < 0.05$ ) different from ADV men; <sup>c</sup> = Significantly ( $p < 0.05$ ) different from REC women; <sup>d</sup> = Significantly ( $p < 0.05$ ) different from REC men; <sup>e</sup> = Significantly ( $p < 0.05$ ) different from CON women; <sup>f</sup> = Significantly ( $p < 0.05$ ) different from CON men.

Significant group effects were found for muscle thickness (VL and BB), pennation angle (BB and TB), fascicle length of TB, and CSA (VM and VL). Compared to CON, ADV possessed greater muscle thickness in VL (*p* = 0.013, BF_10_ = 3.0) and in BB (*p* = 0.012, BF_10_ = 2.2), larger BB pennation angle (*p* = 0.007, BF_10_ = 21.9), and greater CSA in VM (*p* = 0.050, BF_10_ = 2.1) and VL (*p* = 0.009, BF_10_ = 0.7). Compared to REC, ADV possessed greater muscle thickness in VL (*p* = 0.026, BF_10_ = 1.9), larger pennation angle in TB (*p* = 0.009, BF_10_ = 3.2), longer fascicles in TB (*p* = 0.019, BF_10_ = 3.9), and greater CSA in VL (*p* = 0.009, BF_10_ = 0.8); a tendency for greater muscle thickness in BB was also noted for ADV compared to REC (*p* = 0.086, BF_10_ = 0.9). No differences were seen between REC and CON. Morphological comparisons are presented in Table 1.

### Peak aerobic capacity

No significant group x sex interactions were observed for VO_2_peak (F = 1.09, *p* = 0.358, BF_10_ = 10.1), RCT (F = 0.32, *p* = 0.730, BF_10_ = 1.7), or GET (F = 0.05, *p* = 0.949, BF_10_ = 1.1). However, *moderate*-to-*strong* evidence were found in favor of main group effects for each variable. VO_2_peak (F = 9.10, *p* = 0.002, BF_10_ = 17.0) and RCT (F = 5.56, *p* = 0.014, BF_10_ = 4.5) were significantly greater in ADV compared to REC (p ≤ 0.039) and CON (*p* ≤ 0.020), while GET (F = 5.29, *p* = 0.016, BF_10_ = 5.7) was significantly greater in ADV compared to CON (*p* = 0.016) and tended to be greater compared to REC (*p* = 0.087). No differences were seen between REC and CON. Further, the percentage of VO_2_peak for GET and RCT were similar between ADV (GET = 55.2 ± 11.2%; RCT = 71.7 ± 7.5%), REC (GET = 55.9 ± 6.8%; RCT = 73.5 ± 5.9%), and CON (GET = 53.9 ± 4.3%; RCT = 74.6 ± 7.7%). Group differences in absolute measures of aerobic performance are illustrated in Fig 2.

**Fig 2.**
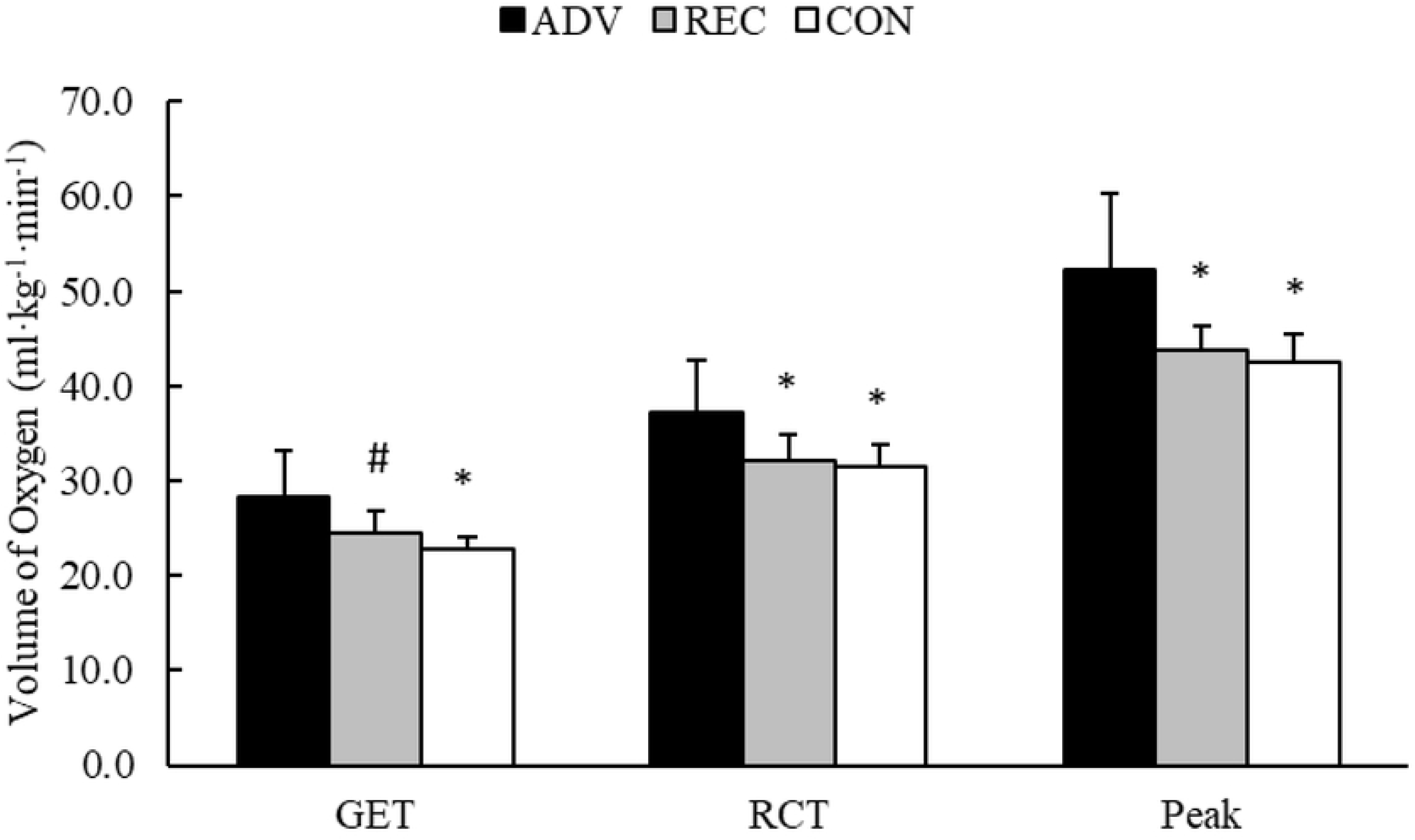
Group differences in aerobic performance measures. *Note*: * = Significantly (p < 0.05) different from ADV. # = Different (p < 0.10) from ADV.

### Resting metabolic rate

Neither a group x sex interaction (F = 0.21, *p* = 0.817, BF_10_ = 0.2) or main group effect (F = 1.67, *p* = 0.220, BF_10_ = 0.1) was observed for RMR recordings in ADV (1788 ± 232 kcal·day^-1^), REC (1768 ± 407 kcal·day^-1^), and CON (1572 ± 356 kcal·day^-1^).

### Body composition

No significant group x sex interactions were observed for any measure of body composition (presented in Table 2). However, the evidence was *strongly*-to-*extremely* in favor of main group effects for body density, regional and total BMC, regional and total lean mass, and BF%. Compared to the REC, ADV possessed greater body density (*p* = 0.004), greater BMC of the arms (*p* = 0.009), greater lean mass (i.e., total and regional; *p* ≤ 0.035), lower BF% (*p* = 0.009), and tended to possess more BMC (total-body: *p* = 0.066; legs: *p* = 0.060) and less fat mass (*p* = 0.064). Compared to CON, ADV possessed greater body density (*p* = 0.006), greater BMC throughout the body (*p* ≤ 0.024), lean mass throughout the body (*p* ≤ 0.009), and lower BF% (*p* = 0.023). No differences were observed between REC and CON.

**Table 2.** Group differences in measures of body composition.

|  | ADV |  |  | REC |  |  | CON |  |  | Group |  | Group x Sex |  |
| --- | --- | --- | --- | --- | --- | --- | --- | --- | --- | --- | --- | --- | --- |
|  | Women | Men | Total | Women | Men | Total | Women | Men | Total | p | BF<sub>10</sub> | p | BF<sub>10</sub> |
| *Anthropometric* |  |  |  |  |  |  |  |  |  |  |  |  |  |
| Height (cm) | 160 ± 13 | 177 ± 3 | 170 ± 11 | 161 ± 4 | 183 ± 8 | 172 ± 14 | 158 ± 4 | 180 ± 9 | 171 ± 14 | 0.785 | >100 | 0.526 | 0.3 |
| Weight (kg) | 68.3 ± 5.0 | 91.5 ± 5.1 | 79.8 ± 13.3 | 59.0 ± 2.0 | 93.5 ± 9.5 | 76.3 ± 19.5 | 60.8 ± 6.3 | 84.9 ± 7.3 | 74.5 ± 14.3 | 0.127 | >100 | 0.169 | 0.3 |
| BMI (kg·m<sup>-2</sup>) | 26.0 ± 3.5 | 29.2 ± 1.9 | 27.6 ± 3.1 | 22.9 ± 1.2 | 28.0 ± 3.4 | 25.5 ± 3.6 | 24.4 ± 3.4 | 26.1 ± 0.7 | 25.4 ± 2.2 | 0.163 | 6.6 | 0.456 | 0.6 |
| Density (kg·L<sup>-1</sup>) | 1.07 ± 0.01 | 1.07 ± 0.01 | 1.07 ± 0.01 | 1.05 ± 0.01 | 1.06 ± 0.01 | 1.05 ± 0.01* | 1.04 ± 0.02 | 1.06 ± 0.01 | 1.05 ± 0.02* | 0.002 | 13.8 | 0.159 | 1.1 |
| *Bone Mineral Content (kg)* |  |  |  |  |  |  |  |  |  |  |  |  |  |
| Total | 3.05 ± 0.38 | 3.75 ± 0.13 | 3.45 ± 0.44 | 2.42 ± 0.16 | 3.62 ± 0.41 | 3.02 ± 0.70# | 2.43 ± 0.14 | 3.29 ± 0.41 | 2.92 ± 0.55* | 0.012 | >100 | 0.299 | 0.7 |
| Arms | 0.45 ± 0.07 | 0.62 ± 0.05 | 0.55 ± 0.11 | 0.32 ± 0.03 | 0.57 ± 0.05 | 0.44 ± 0.14* | 0.30 ± 0.01 | 0.48 ± 0.06 | 0.40 ± 0.11* | 0.001 | >100 | 0.266 | 1.2 |
| Legs | 1.12 ± 0.13 | 1.44 ± 0.11 | 1.30 ± 0.20 | 0.82 ± 0.05 | 1.38 ± 0.17 | 1.10 ± 0.32# | 0.81 ± 0.03 | 1.31 ± 0.22 | 1.09 ± 0.31* | 0.022 | >100 | 0.255 | 0.5 |
| Trunk | 0.95 ± 0.11 | 1.16 ± 0.03 | 1.07 ± 0.13 | 0.79 ± 0.11 | 1.11 ± 0.14 | 0.95 ± 0.21 | 0.82 ± 0.08 | 0.97 ± 0.09 | 0.90 ± 0.11* | 0.028 | >100 | 0.271 | 0.8 |
| *Non-bone fat-free mass (kg)* |  |  |  |  |  |  |  |  |  |  |  |  |  |
| Arms | 7.15 ± 0.89 | 11.12 ± 1.22 | 9.42 ± 2.35 | 4.87 ± 0.49 | 10.02 ± 0.56 | 7.45 ± 2.79* | 4.83 ± 0.42 | 9.04 ± 1.14 | 7.24 ± 2.40* | 0.001 | >100 | 0.400 | 0.6 |
| Legs | 18.4 ± 1.4 | 25.4 ± 1.6 | 22.4 ± 4.0 | 14.3 ± 1.0 | 24.4 ± 1.1 | 19.3 ± 5.4* | 14.7 ± 0.7 | 22.5 ± 3.2 | 19.2 ± 4.7* | 0.008 | >100 | 0.252 | 0.5 |
| Trunk | 27.7 ± 2.9 | 35.2 ± 2.0 | 32.0 ± 4.5 | 20.3 ± 1.2 | 33.5 ± 1.4 | 26.9 ± 7.2* | 21.4 ± 2.2 | 30.1 ± 3.7 | 26.4 ± 5.5* | 0.001 | >100 | 0.073 | 0.8 |
| *4-compartment model* |  |  |  |  |  |  |  |  |  |  |  |  |  |
| Body fat percentage (%) | 11.9 ± 2.4 | 11.0 ± 2.6 | 11.4 ± 2.3 | 23.3 ± 2.4 | 16.1 ± 6.2 | 19.7 ± 5.8* | 23.9 ± 8.4 | 13.7 ± 3.2 | 18.1 ± 7.6* | 0.007 | 16.1 | 0.183 | 2.7 |
| Fat-free mass (kg) | 60.2 ± 3.5 | 81.3 ± 3.4 | 72.3 ± 11.7 | 45.2 ± 2.3 | 78.1 ± 5.3 | 61.7 ± 18.0* | 45.9 ± 1.6 | 73.3 ± 8.4 | 61.6 ± 15.9* | 0.001 | >100 | 0.097 | 0.5 |
| Fat mass (kg) | 8.2 ± 2.1 | 10.2 ± 2.7 | 9.3 ± 2.5 | 13.8 ± 1.4 | 15.4 ± 7.0 | 14.6 ± 4.7# | 14.9 ± 6.7 | 11.5 ± 2.2 | 13.0 ± 4.5 | 0.069 | 1.5 | 0.436 | 0.3 |

### Strength

No significant group x sex interactions were observed for variables obtained from the isometric mid-thigh pull assessment. *Extreme* evidence suggested significant main group effects for F (F = 3.89, *p* = 0.042, BF_10_ = 667,577) and RFD at 200 ms (F = 3.67, *p* = 0.049, BF_10_ = 12,676), as well as tendencies for group differences in F at 150 ms (F = 2.80, *p* = 0.091, BF_10_ = 1,898), F at 200 ms (F = 3.50, *p* = 0.055, BF_10_ = 17,296), F at 250 ms (F = 3.14, *p* = 0.071, BF_10_ = 21524), RFD at 150 ms (F = 2.94, *p* = 0.082, BF_10_ = 1,868), and RFD at 250 ms (F = 3.37, *p* = 0.060, BF_10_ = 20,187). According to post-hoc analysis, ADV produced a higher peak F than CON (*p* = 0.036) and expressed greater RFD at 200 ms than REC (*p* = 0.049). ADV also tended to produce greater F at 200 ms (*p* = 0.062) and 250 ms (*p* = 0.097) compared to REC. No other specific differences were seen between groups. Group differences in F and RFD production across time are illustrated in Fig 3.

**Fig 3.**
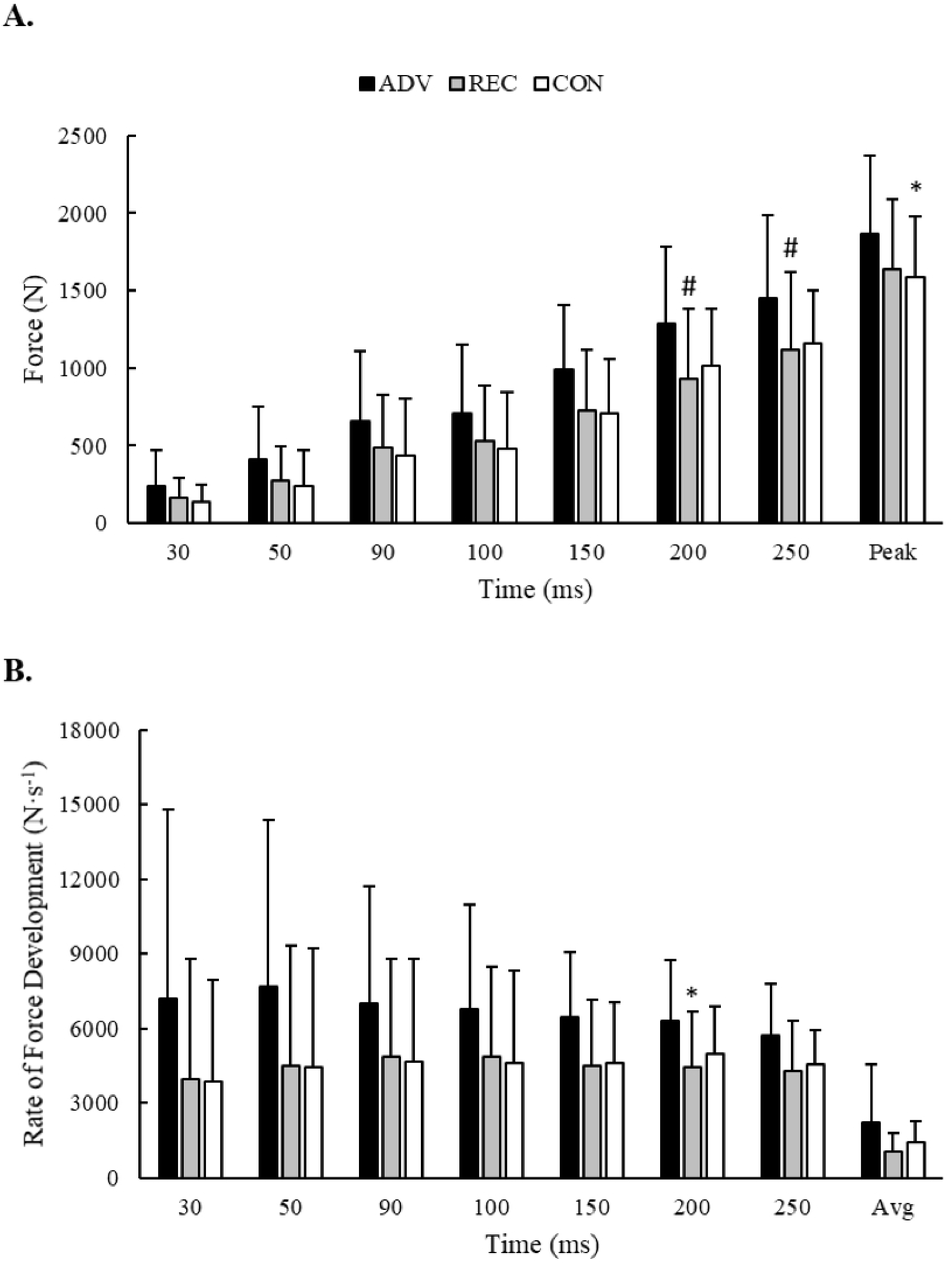
Group differences in A) force and B) rate of force production during an isometric mid-thigh pull. *Note*: * = Significantly (*p* < 0.05) different from ADV. # = Different (*p* < 0.10) from ADV.

### Anaerobic performance

No significant group x sex interactions were observed. *Extreme* evidence in favor of a significant group main effect for CP (F = 7.56, *p* = 0.005, BF_10_ = 267) indicated that ADV possessed a higher CP than REC (*p* = 0.029) and CON (*p* = 0.005). Although extreme evidence was also seen for AWC (F = 4.79, *p* = 0.023, BF_10_ = 247), post-hoc analysis did not reveal specific group differences. No other differences were observed. Group differences in anaerobic performance are illustrated in Fig 4.

**Fig 4.**
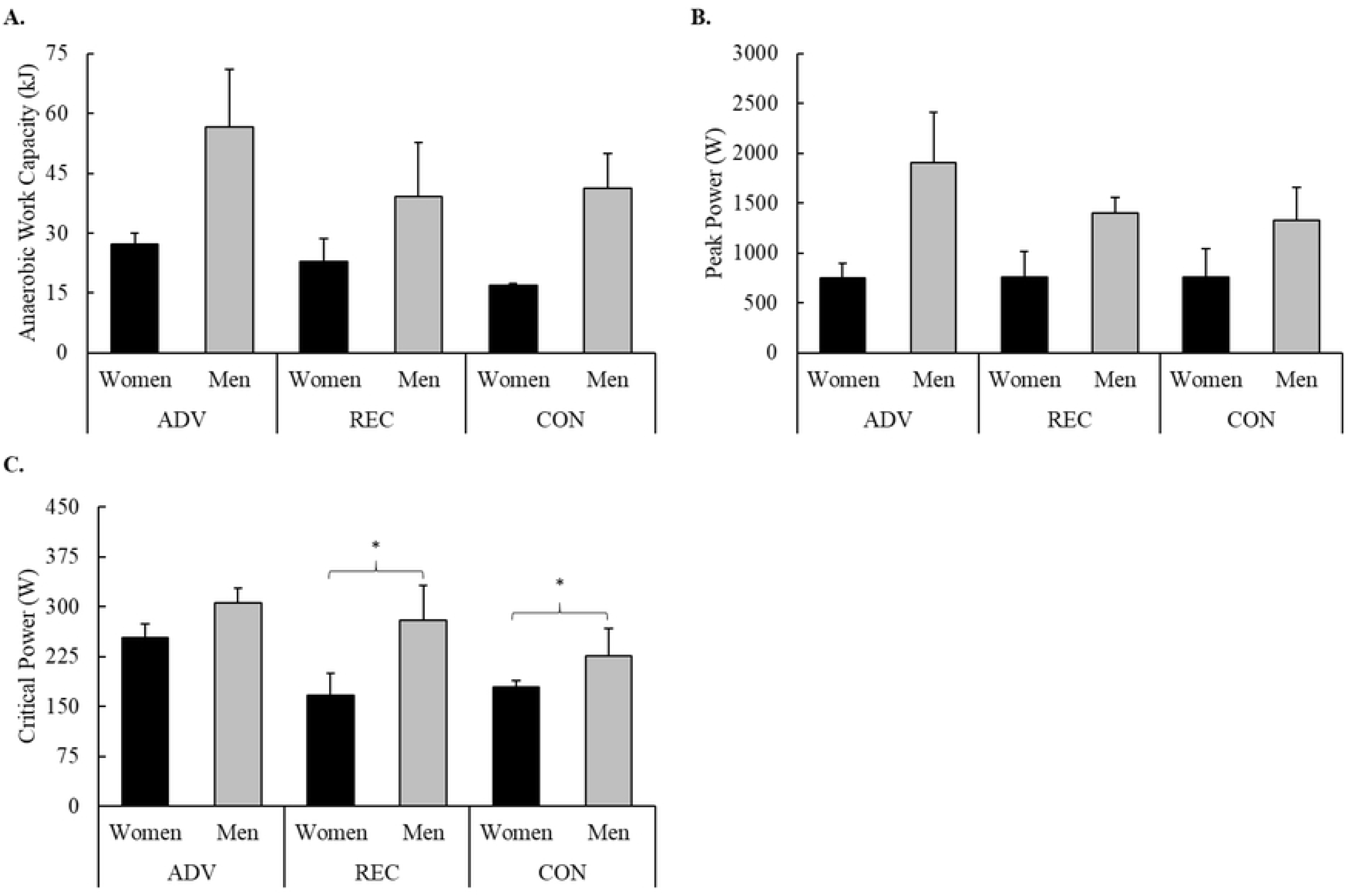
Group differences in A) anaerobic work capacity, B) peak power, and C) critical power. *Note*: * = Significantly (p < 0.05) different than ADV

## DISCUSSION

The primary objectives of this study were to examine anthropometric, hormonal, and physiological differences between advanced CF athletes, recreational CF participants, and resistance and cardiovascular trained adults. Previously, only one other cross-sectional investigation has made physiological comparisons between individuals with at least one year of CF or resistance-training experience (27). The authors reported no differences between the groups except for the CF-trained group possessing greater aerobic ability. This outcome, however, is not surprising considering that the resistance-trained group was not required to also have been performing aerobic exercise. Typical CF training workouts will concurrently incorporate strength and conditioning elements into training (1, 2, 50). Although the conditioning component varies in intensity and duration for each workout, it is important that alternative exercise strategies include both elements to make a fair comparison. The present study builds upon this limitation by having required participants in the CON group to have been participating in both resistance and cardiovascular training on at least 3 days per week each; a similar training frequency was expected of the recreational CF group (i.e., training on at least 3 days per week). Another important aspect of CF training worth consideration is that it includes a wide variety of traditional resistance and aerobic training exercises, along with simple-to-complex gymnastic movements. Proficiency in these movements cannot be assumed after only a year of training and would likely necessitate frequent workout modification. Recently, our group has reported different physiological responses and recovery rates to CF workouts that are completed as prescribed versus those that are modified (i.e., scaled) (11). Thus, CF-trained participants were required to possess at least two years of experience and they were further divided into ADV and REC based upon evidence of their skill as CF athletes (i.e., their previous success in CF competition). Within these contexts, advanced CF athletes were observed to have a more favorable body composition and muscular morphological characteristics, as well as greater aerobic capacity, strength, and ability to sustain high-intensity effort compared to recreational CF participants and physically-active adults. In contrast, no differences were observed between recreational CF participants and physically-active adults in any measure and no differences were seen in resting hormone concentrations or metabolic rate across all groups. This is the first investigation to make comparisons among CF practitioners based on their competitive rank and relative to resistance- and cardiovascular-trained, active adults.

Most competitive CF workouts require athletes to perform 2 or more exercises in a circuit or listwise fashion for several repetitions and rounds, and to do so as quickly as possible or to complete as much work as possible within a given time limit (1, 2, 50). Athletes who can maintain a faster pace or rapidly recover between minimal rest periods would appear to be best positioned to excel in this sport. A recent study in advanced CF athletes, as determined by their performance in a common benchmark workout (i.e., “Fran”), supports this idea (51). Feito et al. (2018) found that the best predictor of repetitions completed during a 15-minute CF workout was the amount of work the athletes could perform on the final trial of four maximal Wingate sprints separated by 90 seconds of rest. In the present study, the ADV group possessed a lower percentage of body fat and greater non-bone fat-free mass compared to the REC and CON groups. In sports, possessing an ideal ratio of skeletal muscle to fat mass may offer a competitive advantage by improving efficiency, thermoregulation, and the ability to sustain effort (52). Aside from their historical success in CF competition, the ADV group’s performance during aerobic and anaerobic testing provide evidence of this ability. ADV participants possessed a higher VO_2_peak than the other groups, which would imply that they were able to perform aerobic work throughout a greater range of workloads (53, 54) but it does not completely explain their ability to sustain effort at higher intensities (55). As the oxygen requirements of a workload exceed an athlete’s capacity to efficiently deliver oxygen, the ability to sustain effort may be further explained by measures of anerobic performance and specific threshold points indicative of the onset of fatigue (i.e., GET, RCT, and CP) (47, 55, 56). Participants in the ADV group were also found to possess a higher GET, RCT, and CP, which are strongly correlated with each other CP (56) and are thought (specifically RCT and CP) to demarcate the point in which exercise transitions from ‘heavy’ to ‘severe’ (56, 57). Together, these data suggest that the ADV athletes in this study had a greater capacity to produce energy aerobically, and that they were better equipped to maintain efforts at higher absolute workloads and thus, be successful in their sport.

Skeletal mass and the morphological characteristics of muscle are suggestive of a greater ability to produce force (58–61). That is, the size, architecture and quality of skeletal muscle reflect the capability of activated muscle to produce force, whereas bone mass provides the structural support and stability needed to effectively translate force production into human movement. In the present study, ADV athletes possessed greater bone and muscle mass/size, larger pennation angles, shorter fascicles, and better quality in the arm and quadriceps musculature compared to the other groups. However, these only partially translated to greater force production by ADV group participants during the IMTP test. IMTP performance was highly variable until 0 to 200 – 250 ms, upon which ADV clearly produced greater force and at a faster rate. The lack of uniformity across all strength measures might be explained by testing specificity and the skillset of our sample. The importance of being able to rapidly activate muscle (i.e., higher RFD) and the magnitude of IMTP force production varies across sports and athletic activities. In weightlifters, significant relationships have been reported between one-repetition maximums in the Olympic lifts and IMTP force (peak and from 0 to 100 – 250 ms) (62) but relationships to RFD have either been limited to specific time bands (from 0 to 200 – 250 ms) (62) or remain unclear in other athletes (63, 64). Although maximal strength in the Olympic and power lifts can distinguish competitive ranking in CF athletes (8, 10), it is not a common requisite of CF competition to maximally perform these lifts. Rather, most competitive workouts either utilize submaximal loads that are performed for several repetitions or they require the athlete to perform maximal (or near maximal) lifts after a fatiguing task (i.e., not a true measure of maximal strength) (50). It is also possible that the composition of the ADV group may help explain the variability observed prior to 200 ms. While all ADV group participants ranked higher than REC in the Open, their participation in later rounds of the Games competition had primarily occurred as part of a team. Within this capacity, team members may be included based on their skill set (e.g., strong/powerful athletes, gymnastically-skilled athletes, endurance athletes) to minimize team weaknesses. This differs from individual competitors who must be proficient in a broader set of skills to be competitive (8, 10). Currently, evidence documenting the physiological differences between high-ranking individual and team competitors does not exist.

There is little evidence to suggest that consistent alterations will occur to resting concentrations in T, C, or IGF-1 as a result of chronic training (14). Rather, their concentrations generally reflect the current status of muscle tissue in response to the demands of training. An overreaching period, marked by elevated training intensity or volume, might elicit transient elevations in T and IGF-1 that typically return to baseline once training returns to ‘normal’ while prolonged overreaching (or overstress) periods may elicit elevations in C (14). CF training is characterized by an effort to maximize training density (i.e., complete a set amount of work as quickly as possible, or maximize work completed within given time frame) within an unplanned (i.e., non-periodized) training structure to promote general physical preparedness (1, 2). Further, the 5-week Open is the most common avenue used by athletes to qualify for the Games (4, 5). Prior to an important competitive event, athletes may elevate training intensity to promote peak performance (65). Thus, the combination of the CF training strategy and the approach of an important, extended competitive event could increase the likelihood of a prolonged period of overstress. The occurrence of which might be identified by changes in resting hormonal concentrations, resting metabolic rate, performance, as well as a variety of other factors (14, 66, 67). However, the present investigation did not reveal any evidence of prolonged stress or negative adaptations to training. Resting hormone concentrations and metabolic rates were similar between groups and the physiological advantages demonstrated by the ADV group appeared to reflect their reported training habits over the past six months (via medical and physical activity history questionnaire). Excluding the conditioning component typically present in CF workouts, members from each group reported using a similar number of sets per muscle group (3 – 6), repetitions (3 – 12), and rest intervals (60 – 90 seconds) during the strength component of their workouts. Only training frequency was reported to be different with the ADV group utilizing a form of resistance exercise on approximately 5.3 days per week whereas the REC and CON groups averaged 4.6 days per week and 3.7 days per week, respectively. Although the greater training frequency seen in ADV would have theoretically provided more of an opportunity to accumulate training volume and promote adaptations, it could have also interfered with their recovery. Nevertheless, ADV possessed a more favorable body composition and generally outperformed the other groups in each performance measure. Therefore, as of one-month prior to competition, adequate recovery appeared to be present in this group. Likewise, the lack of differences seen between REC and CON, who were not actively training for the Open, also provides evidence of adequate recovery. Future investigations can expand on this by monitoring performance surrounding the extended Open competition.

The findings of this study suggest that advanced CF athletes possess a more favorable body composition, greater bone and muscle mass, greater muscle quality and strength, greater aerobic capacity, and a greater ability to sustain effort than recreational CF participants and physically-active adults. The reasons for these differences remain unclear due to the cross-sectional design of this study but may be related to differences in training experience and recent training habits. Although all participants in this study could be considered well-trained (68), ADV group participants reported having more resistance training experience and having been training more frequently over the past 6 months than the other groups. It is possible that their advantages are simply the result of training for a longer amount of time or creating more opportunities to increase their volume load throughout the week. Without documentation (i.e., extensive, detailed training logs), however, it is only possible to speculate upon their potential influence as unknown factors (e.g., training quality, genetic predisposition) would certainly modulate resultant adaptations. Further, the influence of daily variations in the conditioning components of CF workouts on effort and volume load, as well as how these might compare to traditional aerobic exercise (utilized by CON), remains unclear. It is interesting to note that despite the apparent differences in each training strategy (i.e., CF conditioning and traditional aerobic exercise), no differences were seen between REC and CON. To be included in this study, both had to have been regularly participating in their chosen training strategy on 3 – 5 days per week for at least the past year. Nevertheless, REC and CON were found to possess similar physiological characteristics. Future longitudinal investigations that document both the quality and quantity of these training forms may help to provide insight into whether an advantage exists between these strategies or if they promote comparable adaptations among recreationally-active adults.

